# The chromatin-associated PWO proteins interface with circadian and epigenetic signaling to regulate flowering time in Arabidopsis

**DOI:** 10.64898/2026.11.03.763775

**Authors:** Saqlain Haider, Quentin Rivière, Annie Shelton, Ahamed Khan, Alex Webb, Iva Mozgova, Sara Farrona

## Abstract

Flowering time is a major determinant of reproductive success in plants and is regulated by several interconnected pathways which converge on few key flowering time-related genes. In the model plant *Arabidopsis thaliana* (Arabidopsis), both the circadian clock and chromatin-associated proteins integrate environmental cues to coordinate flowering (Staiger *et al*. 2013; Nohales and Kay, 2016; Du *et al*. 2019; Haider and Farrona, 2024). However, how the circadian-epigenetic signaling crosstalk regulates this crucial developmental transition has still been underexplored. PWWP-DOMAIN INTERACTORS OF POLYCOMBS 1 and 2 (PWO1 and PWO2) are chromatin-associated proteins that have been linked to the regulation of histone 2A (H2A) mono-ubiquitination (ub) (H2Aub) levels (Zheng *et al*. 2023). PWO1 and PWO2 also affect flowering time (Hohenstatt *et al*. 2018; Khan *et al*. 2026), but the underlying molecular mechanisms, their effects on clock genes, as well as their distinct contributions to flowering-time regulation remain mostly unknown.

## Introduction

Circadian clocks are endogenous time-keeping systems that generate ∼24-hour biological rhythms and enable organisms to align their physiology with daily and seasonal environmental cues. In Arabidopsis, the circadian oscillator includes two morning expressed MYB transcription factors encoding genes namely *CIRCADIAN CLOCK ASSOCIATED 1* (*CCA1*) and *LATE ELONGATED HYPOCOTYL* (*LHY*), and the evening expressed *TIMING OF CAB EXPRESSION 1* (*TOC1*) gene, also known as *PSEUDO RESPONSE REGULATOR 1* (*PRR1*) (Wang *et al*. 1998; Schaffer *et al*. 1998; Strayer *et al*. 2000; Huang *et al*. 2012). During daytime, the *PRR* gene family is expressed in a sequential order, i.e. *PRR9*, *PPR7*, *PRR5*, *PRR3*, and *TOC1*, each peaking at ∼2hrs interval (Matsushika *et al*. 2000; Nakamichi *et al*. 2010; Nohales and Kay, 2016). The evening complex (EC) comprises *EARLY FLOWERING 3* (*ELF3*), *EARLY FLOWERING 4* (*ELF4*), and *LUX ARRHYTHMO* (*LUX*) (Nohales and Kay, 2016). Additional clock associated proteins such as GIGANTEA (GI), ZEITLUPE (ZTL), and FLAVIN-BINDING, KELCH REPEAT, F BOX 1 (FKF1) interact with core clock components (Imaizumi *et al*. 2003; Mizoguchi *et al*. 2005; Song *et al*. 2012; 14). Altogether, the circadian oscillator regulates numerous physiological and metabolic processes in plants (McClung, 2011; Staiger *et al*. 2013; Nohales and Kay, 2016; Oakenfull and Davis, 2017).

The circadian clock plays a fundamental role in the regulation of flowering time through the so-called photoperiodi pathway. The photoperiodism is the process by which plants measure the changes in day and night cycles associated with the seasons, and make accordingly floral transition (Samach *et al*. 2000; Blackman, 2017). Mechanistically, circadian oscillators regulate photoperiodic response at both transcriptional and pos-transcriptional levels. In Arabidopsis, the circadian clock rhythmically maintains the expression of the key positive flowering gene *CONSTANS* (*CO*) late, at a time that corresponds to the night under non-inductive short day (SD) (Suárez-López *et al*. 2001; Yanovsky and Kay, 2002). CONSTITUTIVE PHOTOMORPHOGENIC1 (COP1) degrades CO in the dark via the proteasomal pathway and the reduced CO activity limits *FLOWERING LOCUS T* (*FT)* (key florigen) induction, delaying flowering (Jang *et al*. 2008). CO keeps being degraded until the photoperiod becomes long enough, and the peak of the CO protein coincides with daylight in long day (LD) (Yanovsky and Kay, 2002, Valverde *et al*. 2004).

The circadian clock is mainly involved in that process through GI and PRRs. GI regulates *CO* mRNA by degrading CYCLING DOF TRANSCRIPTION FACTORS (CDFs) (Fornara *et al*. 2009). CDFs, among which CDF1 and CDF2 play a major role, redundantly repress *CO* transcription by directly binding to its promoter (Imaizumi *et al*. 2005; Imaizumi *et al*. 2003; Fornara *et al*. 2009; Hayama *et al*. 2017) and can also directly repress *FT* expression (Corbesier *et al*. 2007; Jaeger and Wigge, 2007; Song *et al*. 2012). In addition, PRRs, including PRR9, PRR7, and PRR5, also directly repress *CDFs* (primarily *CDF1*) transcription thus promoting *CO* expression (Nakamichi *et al*. 2007). At post-transcriptional level, GI contributes to CO protein stability at the end of the day in LD (Song *et al*. 2014; Hwang *et al*. 2019) and PRRs physically interact with CO to prevent its degradation by COP1 (Hayama *et al*. 2017). On the other hand, FLOWERING LOCUS C (FLC) represses *FT* transcription by directly binding to its promoter, thus preventing precocious flowering (Searle *et al*. 2006).

Here, we conducted extensive phenotyping and data mining and found that PWOs play a decisive role in the photoperiodic pathway in concert with the circadian clock.

### PWO2 has a stronger effect on flowering time than PWO1

To better understand the interaction between PWO1 and PWO2 in flowering time regulation, we scored the flowering time phenotypes of the *pwo1-1* (*pwo1)*, *pwo2-2* (*pwo2*), and *pwo1-1;pwo2-2* (*pwo1;2*) mutants versus Col-0 plants under LD conditions (Figure 1a). The *pwo1* mutants had a weak flowering phenotype in terms of leaf number in line with a previous study (Hohenstatt *et al*. 2018) (Figure 1b). However, *pwo1* mutant was similar to Col-0 in terms of number of days (Figure 1c). Contrarily, the *pwo2* mutant was considerably delayed compared to Col-0 *i.e.,* higher number of leaves alongside number of days (Figure 1b,c). Notably, the flowering phenotype of *pwo1;2* mostly resembled that of *pwo2* both in terms of days to flower and number of leaves (Figure 1b-c). A downward trend in terms of number of leaves in *pwo1;*2 in comparison with *pwo2* was consistent with an additive effect of the two mutations, but the absence of statistical difference prevented us from reaching a definitive conclusion. Nevertheless, our data clearly indicated that flowering is primarily controlled by PWO2. We also analyzed the flowering phenotype of Col-0 vs *pwo1;2* under SD conditions which revealed a slow growth trend in *pwo1;2* (Supplementary figure 1a). Interestingly under SD, the *pwo1;2* double mutant produced lesser number of leaves compared to Col-0 plants despite being delayed in terms of number of days (Supplementary figure 1b-d).

**Figure 1.**
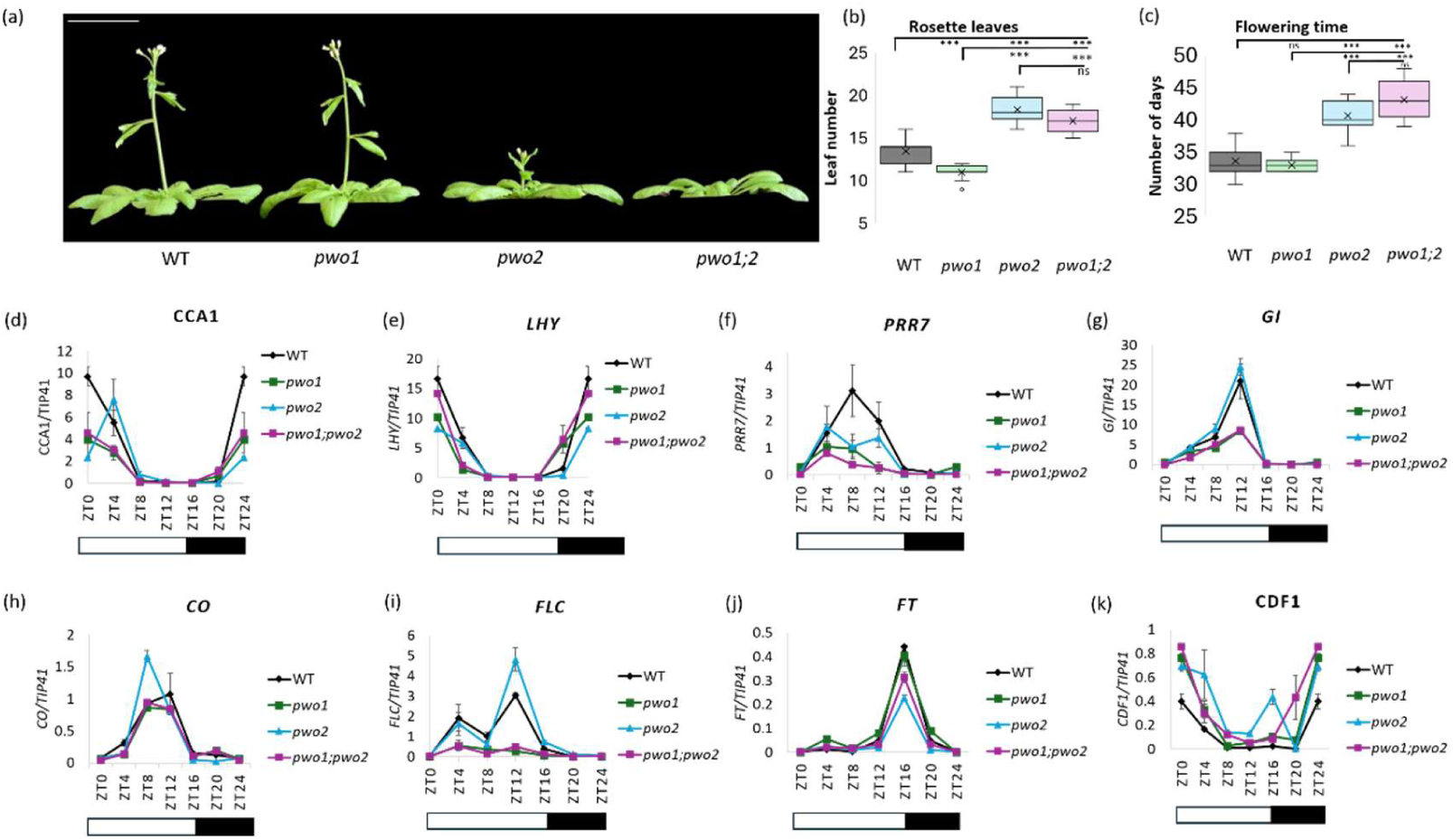
*PWO1*/*2* regulate flowering time and the expression of clock and photoperiod genes under LD conditions. (a) Representative images of 40-day-old Col-0 (WT), *pwo1*, *pwo2* and *pwo1;2* plants. Scale bar = 5cm. (b) Rosette leaf number of Col-0, *pwo1*, *pwo2* and *pwo1;2* double mutants grown under LD conditions. (c) Number of days till flowering for Col-0, *pwo1*, *pwo2* and *pwo1;2* plants grown under LD conditions. At least 10 plants per genotype were scored. (d-K) Expression levels of (d) *CCA1*, (e) LHY, (f) *PRR7*, (g) *GI*, (h) *CO*, (i) *FLC*, (j) *FT* and (k) *CDF1* analyzed in 10-day-old Col-0, *pwo1*, *pwo2*, *pwo1;2* seedlings grown under LD conditions harvested every 4-hours. White and black bars represent day and night times respectively. *TIP41* was used as an internal control. At least 3 biological replicates were used.

### *PWO1/2* regulate diel expression of flowering regulator genes

To understand the impact of *PWO1/2* on flowering regulators, we first examined the expression profiles of *FLC* and *FT* (key flowering regulators) in Col-0 vs *pwo1;2* from 6-day-old and 10-day-old seedlings harvested at ZT16. Both *FLC* and *FT* were reduced in *pwo1;2* regardless of the developmental age of the plants (Supplementary figure 2). A reduced *FT* despite lower *FLC* indicated a possible defect in the photoperiod pathway. To this end, we first analyzed the binding of PWO1-FLAG using ChIP-seq data from Zheng *et al*. (2023) which indicated PWO1-FLAG binding to *TOC1*, *PRR7*, *PRR5*, *ELF3*, and *GI* (Supplementary figure 3). Expression analyses of these genes revealed a reduction in the expression of *PRR7* and a non-significant decline in the other tested genes in *pwo1;2* (Supplementary figure 4). As core clock components have been implicated in regulation of photoperiodic flowering (Fowler *et al*. 1999; Mizoguchi *et al*. 2005; Nakamichi *et al*. 5; 2007; Song *et al*. 2014), we hypothesized PWO1/2 may control flowering time through regulation of circadian clock and photoperiod pathway genes.

To investigate the possible role of PWOs in regulating the diel oscillations of key clock and flowering genes, we harvested samples from 10-day-old long day (LD)-grown seedlings every 4 hours from ZT0 to ZT24 (Figure 1d-k). To this end, we examined the expression profiles of the *CCA1*, *LHY*, *PRR7*, *GI*, *CO*, *FLC*, *FT*, and *CDF1* genes with a well-established role in control of the photoperiod pathway (Schaffer *et al*. 1998; Suárez-López *et al*. 2001; Mizoguchi *et al*. 2005; Searle *et al*. 2006; Corbesier *et al*. 2007; Nakamichi *et al*. 2007; Hayama *et al*. 2017).

*CCA1* had a reduced expression at ZT0 in *pwo1*, *pwo2*, and *pwo1;2* mutants, although the rhythmic profile was maintained across a diel cycle (Figure 1d). *LHY* expression was similar to Col-0 in *pwo1;2* while it was visibly reduced in *pwo1* and *pwo2* at ZT0 (Figure 1e). *PRR7* expression was altered in all *pwo* mutant backgrounds. In *pwo1* mutants, *PRR7* transcript levels were reduced from ZT8 onwards. In *pwo2* mutants, *PRR7* had an altered pattern with peaks at ZT4 and ZT12, although expression was strongly reduced at ZT8. Notably, *pwo1;2* mutants had a near-complete loss of *PRR7* rhythms together with near loss of expression (Figure 1f). For *GI,* a pronounced reduction at ZT12 was observed in *pwo1* and *pwo1;2* mutants while *pwo2* had WT-like expression (Figure 1g). *CO* expression remained largely unaffected in all *pwo* mutants across the diel cycle (Figure 1h). *FLC* expression was not rhythmic in both *pwo1* and *pwo1;2* mutants. By contrast, *pwo2* mutants maintained a largely WT-like expression pattern, although a stronger peak was observed at ZT12 (Figure 1i). In *pwo1* mutants, *FT* expression was transiently elevated at ZT4 possibly explaining the slight early flowering of *pwo1* mutant as a *FT* expression peak at ZT4 is sufficient for floral induction under natural light conditions (Song *et al*. 2018). In *pwo2* mutants, *FT* expression was strongly reduced at ZT16 in line with the delayed flowering phenotype while *pwo1;2* also had reduced expression at ZT16 (Figure 1j). Finally, all the *pwo* mutants had a higher *CDF1* expression at ZT0 with *pwo2* uniquely showing a strong upregulation at ZT16 and *pwo1;2* also showing higher *CDF1* expression at ZT20 (Figure 1k). Taking together, these results indicate that the diel oscillation patterns of key flowering regulatory genes are disrupted in *pwo1*, *pwo2*, and *pwo1;2* mutants.

The impact of PWO1/2 on rhythms of the circadian oscillator gene *PRR7* prompted us to investigate whether these mutants also affect the free running circadian period. We estimated the free-running period in wild-type and *pwo1, pwo2, pwo1;2* mutant plants in constant light and temperature using leaf movement (Herrero *et al*. 2026). However, no significant differences were detected between these genotypes and Col-0, indicating that loss of *PWO1/2* function does not significantly alter circadian period under the conditions examined (Supplementary figure 5). These findings are also supported by a similar WT-like trend for *CCA1* and *LHY* genes in *pwo1; pwo2*; *pwo1;2* mutants (Figure 1d-e). Nevertheless, as in our diel expression analysis, two key genes regulating flowering, *PRR7* and *GI*, showed strong perturbations in *pwo1;2* compared to Col-0 (Figure 1f-g), we further tested whether PWO1/2 may control flowering via *PRR7* and *GI* transcriptional regulation.

### *PRR7* is epistatic to *PWO1/2* for flowering time control

To better understand the genetic relationship between *PWO1/2* and *PRR7* for flowering time control, we crossed *pwo1;2* with *prr7-11 (prr7)* to obtain the *pwo1;2;prr7* triple mutant. Flowering time analysis of *pwo1;2;prr7* triple mutant and *prr7* revealed an epistatic relationship between *PWO1/2* and *PRR7* both in terms of number of days and rosette leaves (Figure 2a). Briefly, *pwo1;2* and *prr7* flowered at 42 days and 59 days, respectively, while flowering in *pwo1;2;prr7* occurred at 60 days. On the other hand, *pwo1;2* produced 17 leaves, *prr7* 40 leaves, and *pwo1;pwo2;prr7* 38 leaves at flowering (Figures 2b-c). Such results were in line with an epistatic relationship between *PRR7* and *PWO1/2* for the control of flowering time.

**Figure 2.**
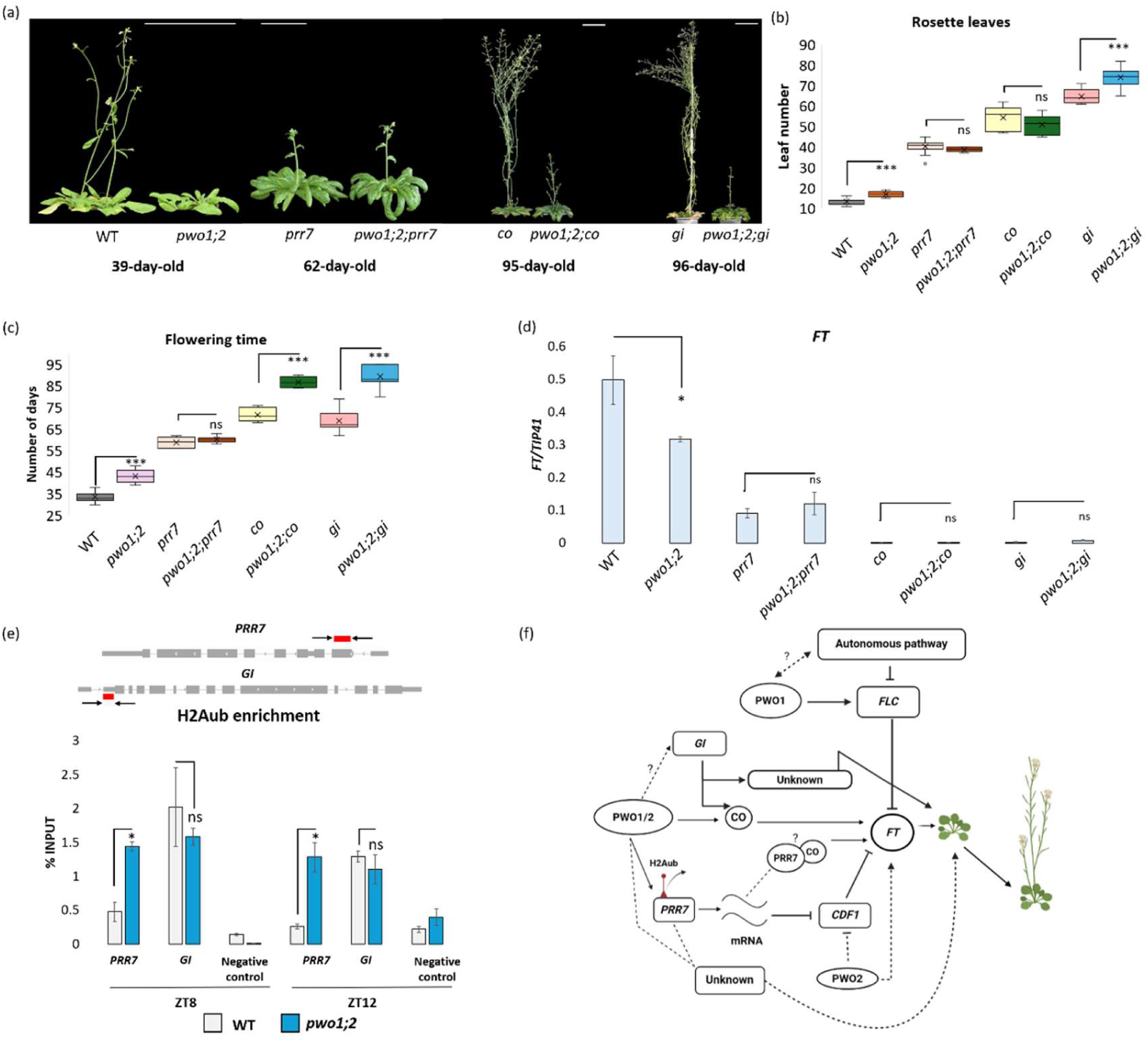
*PWO1/2* modulates flowering time through the regulation of *PRR7*. (a) Representative images of 39-day-old WT and *pwo1;2* plants, 62-day-old *prr7*, and *pwo1;2;prr7* mutants, 96-day-old *gi* and *pwo1;2;gi* mutants, and 95-day-old *co*, and *pwo1;2;co* triple mutants. Scale bars = 5cm. (b) Rosette leaf number of Col-0, *pwo1;2, prr7*, *pwo1;2;prr7*, *gi*, *pwo1;2;gi*, *co,* and *pwo1;2;co* mutants at the time of flowering. (c) Number of days till flowering for Col-0 *pwo1;2, prr7*, *pwo1;2;prr7*, *gi*, *pwo1;2;gi*, *co,* and *pwo1;2;co* mutants grown under LD conditions. (d) Expression analysis of *FT* in 10-day-old seedlings in Col-0 *pwo1;2, prr7*, *pwo1;2;prr7*, *co, pwo1;2;co gi,* and *pwo1;2,gi* mutants. Samples were harvested at the end of long day (ZT16). Error bars represent SD of three biological replicates. Student t-test was used to calculate the significance level. (e) Primers spanning the TSS of *PRR7* and proximal promoter of *GI* locus. Red bar: analyzed region. (f) ChIP-assays from samples harvested at ZT8 and ZT12. Increased H2Aub levels downstream of *PRR7* TSS was detected. An intergenic region (*AG.i*) which was previously shown to harbor no H2Aub enrichment was selected as negative control (Godwin *et al*. 2024). ChIP-qPCR results shown as % of input DNA. Student t-test was used to calculate the significance level. Two biological and three technical replicates were used. (g) A model for the role of *PWO1/2* in control of flowering time through *PRR7-CO* and an additional unknown pathway. In this model, PWOs regulate the expression of *PRR7* via H2Aub removal. Once *PRR7* mRNA is synthesized and translated, the PRR7 protein may interact with CO to prevent its degradation and promote flowering through the *CO-FT* route (Nakamichi *et al*. 2007; Huang *et al*. 2017). Although further work is required to confirm CO protein activity and abundance in *pwo1;2* mutant. PRR7 mediated *CDF1* repression is essential to promote flowering under LDs (Nakamichi *et al*. 2007). Here, PWO2 may uniquely promote *FT* expression by repressing *CDF1* expression (in a PRR7 associated pathway), thus limiting CDF1-mediated *FT* repression (Song *et al*. 2012). On the other hand, an additive interaction for leaves and synergistic interaction for days between *PWO1/2-GI* indicates that the proteins possibly work in parallel and may converge on similar pathway(s), although independent of the *CO-FT* branch, possibly through age- or growth-related pathways. PWO1 may interact with autonomous pathway genes to regulate *FLC* expression (Hohenstatt *et al*. 2018), providing an additional layer of flowering time regulation. Solid lines: interaction known or proved through this study. Dashed lines: Proposed interactions.

We next wanted to clarify whether the observed epistatic interaction between *PWO1/2* and *PRR7* converges on *CO* for flowering time control (Nakamichi *et al*. 2007). To this end, we crossed the *pwo1;2* double mutant with the *co-101* (*co*) single mutant to obtain triple *pwo1;2;co* mutant plants (Figure 2a). While *co* plants produced ∼51 leaves, *pwo1;2;co* plants produced ∼48 leaves at the time of flowering (Figure 2b). These results confirmed that *CO* is epistatic to *PWO1/2* for flowering in terms of leaf number. Nevertheless, *pwo1;2;co* triple mutant was significantly delayed when compared to *co* considering days to flower: *co* flowered on average on day 69, while the *pwo1;2;co* triple mutant flowered on day 84 (Figure 2c). Therefore, *CO* and *PWO1/2* showed a synergistic interaction for flowering in terms of number of days.

As *GI* is a dominant regulator of *CO* expression and protein stability under LDs, we crossed *pwo1;2* double mutants with *gi-2 (gi)* to obtain the *pwo1;2;gi* triple mutant (Figure 2a). The flowering phenotype revealed that *gi* and *pwo1;2* mutants flowered ∼36 days and ∼9 days later than Col-0 respectively (Figure 2c). On the other hand, *pwo1;2;gi* mutants flowered ∼58 days later than Col-0, indicating that the *pwo1;2* genetic background enhanced the phenotype of *gi* (Figure 2c). While for rosette leaf number, *pwo1;2* and *gi* produced ∼4 and ∼50 extra leaves compared to Col-0, *pwo1;2;gi* produced ∼60 extra leaves (Figure 2b), indicating an additive or possible close to synergistic genetic interaction between *gi* and *pwo1;pwo2* for rosette leaf number.

To determine whether the epistatic interactions between *PWO1/PWO2* and *PRR7* or *CO* converge on the expression of *FT*, *FT* transcript levels were quantified in the Col-0, *pwo1;2*, *prr7*, *pwo1;2;prr7*, *co*, *pwo1;2;co*, *gi*, and *pwo1;2;gi* genotypes from samples harvested at ZT16. Compared to WT, *prr7* showed reduced *FT* expression as reported previously (Nakamichi *et al*. 2007), while it remained the same as *prr7* in *pwo1;2;prr7,* in line with a similar flowering phenotype. A complete loss of *FT* expression was observed in *co* and *gi* (Figure 2d), as expected from previous reports (Suárez-López *et al*. 2001; Mizoguchi *et al*. 2005). Similar to *co* and *gi*, low *FT* mRNA levels were also quantified in the *pwo1;2;co* and *pwo1;2;gi* mutant (Figure 2d).

### PWO1/PWO2 regulate H2Aub levels at the *PRR7* locus

Polycomb repressive complex 1 (PRC1) deposits the repressive H2Aub mark at its target genes (Bratzel *et al*. 2010). On the other hand, PWO1/2 facilitate the removal of the H2Aub mark by UBIQUITIN SPECIFIC PROTEASE 5 (UBP5) (Zheng *et al*. 2023; Godwin *et al*. 2024). Therefore, considering our expression and phenotypic data, we sought to clarify whether both *PRR7* and *GI* show variation in H2Aub levels between WT and *pwo1;2* (Figure 2e). We harvested samples from 10-day-old seedlings at ZT8 and ZT12, when *PRR7* and *GI* expression reached their peak in WT plants and showed drastically reduced expression in *pwo1;2* (Figure 1f-g). A significant increase in H2Aub enrichment was observed in *pwo1;2* compared to Col-0 just downstream of *PRR7* TSS at both timepoints (ZT8, ZT12) (Figure 2e). However, we did not observe any changes for *GI* between WT and *pwo1;2*. Despite low *GI* expression (Figure 1g), we could not detect a change in H2Aub levels between WT and *pwo1;2* (Figure 2e). Indeed, data mining from Godwin *et al*. (2024) confirmed binding of UBP5 to *PRR7* locus but not *GI*, despite a possible binding of PWO1 to *GI* (Supplementary figure 3; Zheng *et al*. 2023).

### Do *PWO1/2* also regulate flowering independently of *CO*-dependent photoperiod pathway?

A central finding from our work is the fact that PWOs regulate the CO-dependent photoperiod pathway through the transcriptional regulation of *PRR7* at the epigenetic level. However, the photoperiod pathway is probably not the only flowering pathway impacted by PWOs, as suggested by the synergistic effects of *CO* and *PWO1/2* in the number of days for flowering (Figure 2c). This hypothesis is further strengthened by the additive and synergistic interaction between *PWO1/2* and *GI* for rosette leaf number and for number of days despite similar *FT* levels. The potential additional flowering pathway(s) influenced by PWOs were likely disrupted in the *prr7* loss-of-function background, explaining why *PRR7* appeared epistatic to *PWO1/2* (Figure 2a) despite though *PRR7* reported to regulate flowering time through *CO* (Figure 2b) (Nakamichi *et al*. 2007). With such consideration, our results did not contradict the previous observation by Hohenstatt et al. (2018) that *PWO1* interacts with *FLC* to control the floral transition. This prompted us to evaluate another circadian clock gene, which uses both CO and alternative routes to regulate the flowering time, *ELF3* (Kim *et al*. 2005). We observed a misregulation of *ELF3* expression in all *pwo* mutants (Supplementary figure 6), providing new avenues for investigating the interplay between PWOs, the circadian clock and the floral transition.

One additional candidate pathway that PWOs may interact with is the age-related one. The *pwo1;2* mutants grown under SD showed slower growth and leaf production rate (number of leaves at flowering/number of days at flowering), with Col-0 showing 0.79 leaves/days, while *pwo1;2* having a rate of 0.45 leaves/days (Supplementary figure 1a). A non-significant, yet mild delay in juvenile-to-adult transition was also observed under LD conditions in *pwo1;2* (Supplementary figure 7). Interestingly, under LD conditions, introduction of *pwo1;2* into the *gi* background slows the growth and delays the leaf production of the *gi* mutant (Supplementary figure 8).

A non-photoperiodic role of *PWO1/2* is also suggested by an early flowering phenotype of *pwo1;2* under SDs (Supplementary figure 1b,c). A complete reduction of *FLC* expression in *pwo1* and a similar trend in *pwo1;2* but not in *pwo2* (Figure 1i) also pointed out to a possible interaction specifically between PWO1 and the autonomous pathway (Figure 2f) in which *FLC* is involved (Rouse *et al*. 2002; Qi *et al*. 2022). Interestingly, the expression of *FLC* returned to WT levels in *pwo1;2;prr7* and *prr7* (Supplementary figure 9). These results indicate the broader impact of PWO1/2 on PRR7-associated pathways while also revealing the distinctive interactions between PWO1/2 and circadian and photoperiod pathway genes.

## Material and Methods

### Plant material and growth conditions

*Arabidopsis thaliana* accession Columbia-0 (Col-0) was used as the wild type (WT) in this study. All the mutant genotypes used in this study were in Col-0 background; these include *pwo1-1* (*pwo1*), *pwo2-2* (*pwo2*), *pwo1-1;pwo2-2* (*pwo1;2*), *gi-2* (*gi*), *co-101* (*co*), and *prr7-11* (*prr7*). These mutants have been previously described (Hohenstatt *et al*. 2018; Khan *et al*. 2026; Fowler *et al*. 1999; Takada *et al*. 2003; Nakamichi *et al*. 2005; 7). The triple mutants of *pwo1;pwo2;gi*, *pwo1;pwo2;co*, and *pwo1;pwo2;prr7* were obtained by crossing *pwo1;2* with the single mutants of *gi*, *co*, *prr7*. Homozygous F3 lines were further used for subsequent analysis. Primers used to confirm the mutant genotypes are mentioned in supplementary table 1.

Plants were grown in pots containing peat, vermiculite, and perlite in a 5:1:1 ratio, in long day (LD) (16/8-light/dark) conditions under fluorescent lamps at 120 μmol m−2 s−1 at a temperature regime of 21°C. For short days (SD) experiments, plants were grown under similar temperature regime with SD (8/16-light/dark) conditions. For *in vitro* growth, seeds were sowed on ½ Murashige & Skoog (MS) medium supplemented with 0.8% agar and 0.5% sucrose. Plates were then stratified at 4°C for 3 days for breaking dormancy and synchronizing germination.

### Flowering time analysis

Flowering time was scored by counting both the number of rosette leaves (excluding cotyledons) and number of days when the first floral bud opened. All the experiments were repeated twice independently, and similar results were obtained (Supplementary table 2-7).

### Statistical analysis

Differences in flowering time and leaf numbers among genotypes were analyzed separately for each experimental replicate using Welch’s one-way ANOVA followed by Games–Howell post-hoc comparisons. Statistical significance was defined as adjusted *p* < 0.05.

### Free running period estimation

Circadian period was measured using leaf movement as described in Herrero *et al*. (2026). Plants were grown in 12L:12D until 3 days after germination (DAG), transferred to LL, and monitored for five more days. The free-running circadian period was estimated from leaf-movement time series using BioDare2 (Zieliński *et al*. 2024). Individual traces were assessed for period, relative amplitude error (RAE), and oscillation amplitude. Traces with poor fits (RAE > 0.5) or estimated periods outside 18–30 h were excluded before genotype means were calculated (Supplementary table 8).

### RNA extraction, cDNA synthesis, and qPCR

RNA was extracted from 25-50 Arabidopsis seedlings for each biological replicate from each line grown on an ½ MS agar plate by Monarch Total RNA Miniprep Kit (NEB) with on column DNase treatment. 1 μg of total RNA was used to synthesize first strand cDNA using RevertAid First strand cDNA synthesis kit by Thermo Scientific. The PCR amplification from cDNA was carried out using Takyon™ No ROX SYBR 2X MasterMix blue dTTP (Eurogentec). Expression analyses were performed by real-time PCR (Roche LightCycler). *TAP42 INTERACTING PROTEIN OF 41 kDa* (*TIP41*) was used as housekeeping gene. Error bars show standard deviation of three biological replicates. Student’s t-test was used to calculate the statistical significance of results. A list of primer sets used to amplify genes is presented in supplementary table 9.

### Chromatin-immunoprecipitation coupled to qPCR (ChIP-qPCR)

ChIP assays were performed as published recently (Khan *et al*. 2026). Briefly, plants were grown for 10 days in LDs and harvested at ZT8 and ZT12. Plant tissues were cross-linked using 1% formaldehyde under vacuum. Chromatin immunoprecipitation of WT and *pwo1;2* was performed by using *anti-H2Aub* antibody (Cell Signalling Technology D27C4). Input DNA was diluted 1:10 and IP by 1:5, and 2 μL of immunoprecipitated DNA was used for quantitative PCR (qPCR). ChIP-qPCR was performed using a LightCycler® 480 Instrument (Roche) with Takyon™ No ROX SYBR 2× MasterMix Blue dTTP (Eurogentec). Primers used to amplify the regions are mentioned in supplementary table 10. Two biological replicates and three technical replicates were used for ChIP-qPCR. Student’s t-test was used to calculate the statistical significance of results.

## Supporting information

Supplementary_Figures.zip

Supplementary_Tables.zip

## Author contributions

SH and SF conceptualized the study, experimental approach, and methodology. SH performed the genetic crossing, genotyping, phenotyping, expression profiling, and ChIP-assays. QR provided critical insights during all stages of manuscript preparation. AS performed the leaf movement assay. AK performed initial genetic analyses with *pwo* mutant lines. SH, QR, and SF analyzed the data. SH and SF wrote the manuscript. QR, IM, and AW reviewed the manuscript.

## Acknowledgments

SF and SH would like to thank Research Ireland (grant number 20/ FFP-P/8693) for funding this work. IM would like to thank the Czech Academy of Sciences (Fellowship *Lumina quaeruntur –* LQ200961901) and QR is grateful to MSCA Fellowship CZ (EUROTALENTS - CZ.02.01.01/00/22_010/0008764) for funding. AS is grateful to the Gates Cambridge Trust for funding. SH and SF would like to thank Professor Rossana Henriques from University College Cork (Ireland) for providing the *gi-2* and *co-101* seeds. The *prr7-11* seeds were kindly provided by Professor Paloma Mas from Center for Research in Agricultural Genomics (Spain).

## Supplementary figure legends

**Supplementary figure 1**. *pwo1;2* mutants are early flowering short day (SD) conditions.

**Supplementary figure 2**. Expression analysis of *FLC* and *FT* in 6- and 10-day-old WT vs *pwo1;2* seedlings.

**Supplementary figure 3**. PWO1-FLAG binding to circadian clock genes.

**Supplementary figure 4**. Expression of selected PWO1 target genes in wild-type and *pwo1;2* plants.

**Supplementary figure 5**. Circadian period length based on rhythmic leaf movement in *pwo1, pwo2, pwo1;2* mutants compared to WT.

**Supplementary figure 6**. Diel expression profiles of *ELF3* in *pwo1*, *pwo2*, *pwo1;2* compared to Col-0.

**Supplementary figure 7**. Abaxial trichome analysis of Col-0 and *pwo1;2* under LD conditions.

**Supplementary figure 8**. Vegetative development and leaf production in *pwo;1;2;gi* compared to *gi* under LD conditions.

**Supplementary figure 9**. *FLC* expression analysis 10-day-old seedlings in Col-0 and mutant genotypes.

## Supplementary table legends

**Supplementary Table 1**. Genotyping primers used in this study to confirm Col-0 and mutant alleles.

**Supplementary table 2.** Raw data and statistical analysis for flowering in terms of number of days (Replicate 1).

**Supplementary table 3.** Raw data and statistical analysis for flowering in terms of number of leaves (Replicate 1).

**Supplementary table 4.** Raw data and statistical analysis for flowering in terms of number of days (Replicate 2).

**Supplementary table 5.** Raw data and statistical analysis for flowering in terms of number of leaves (Replicate 2).

**Supplementary table 6.** Raw data and statistical analysis for flowering in terms of number of leaves under SD (Rep 1 and 2).

**Supplementary table 7.** Raw data and statistical analysis for flowering in terms of number of days under SD (Rep 1 and 2).

**Supplementary table 8.** qPCR primers used in this study to analyze the expression profiles of genes

**Supplementary table 9.** ChIP-qPCR primers used in this study to analyze H2Aub enrichment.

**Supplementary table 10.** Free running circadian period raw values.

