## Supplementary_Figures.zip for "The chromatin-associated PWO proteins interface with circadian and epigenetic signaling to regulate flowering time in Arabidopsis": Haider et al_Supplementary figures.pdf

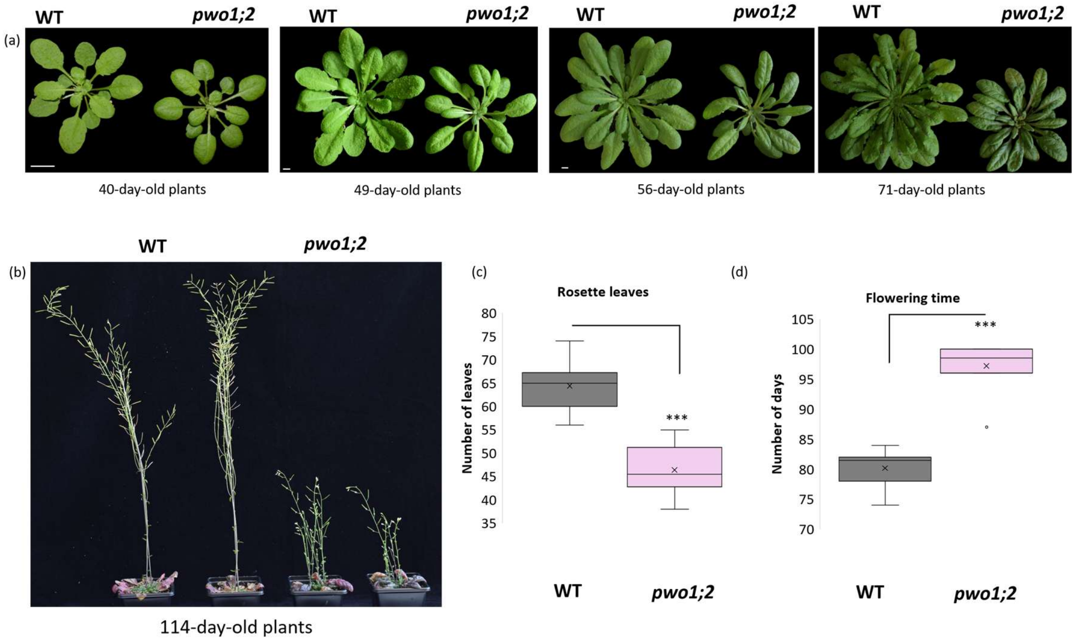


**Supplemenatry figure 1. *pwo1;2* mutants are early flowering under short day (SD) conditions.** (a) Representative images of Col-0 (WT) and *pwo1-1;pwo2-1* (*pwo1;2)* mutants across different timescales. (b) Representative images of 114-day-old WT and *pwo1;2* plants. (c) Rosette leaf number of WT and *pwo1;2* double mutants at flowering time grown under SD conditions. (d) Number of days till flowering for WT and *pwo1;2* double mutants grown under SD conditions. At least 10 plants per genotype were scored. Experiment was repeated and similar results were obtained. Student's t-test was used to calculate the statistical significance of the results. Asterisks indicate significant differences between genotypes; *=P < 0.05, **=P < 0.01, *** = P < 0.001.

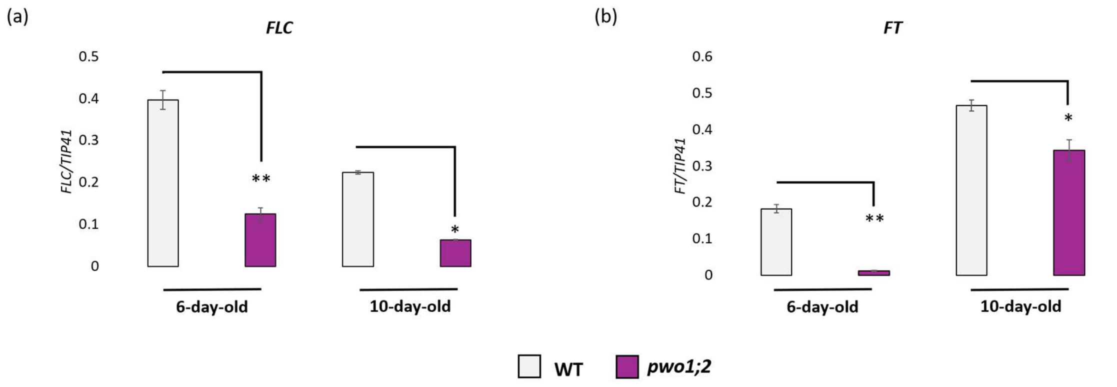


**Supplementary figure 2. Expression analysis of *FLC* and *FT* in 6- and 10-day-old Col-0 (WT) vs *pwo1;2* seedlings.** (a-b) Relative transcript levels of (a) *FLC* and (b) *FT* in 6- and 10-day-old WT and *pwo1;2* seedlings. Transcript levels were normalized to *TIP41*. Error bars represent the standard deviation from two biological replicates. Student's t-test was used to calculate the significance level. Asterisks indicate significant differences between genotypes at the indicated ages; *=P < 0.05, **=P < 0.01.

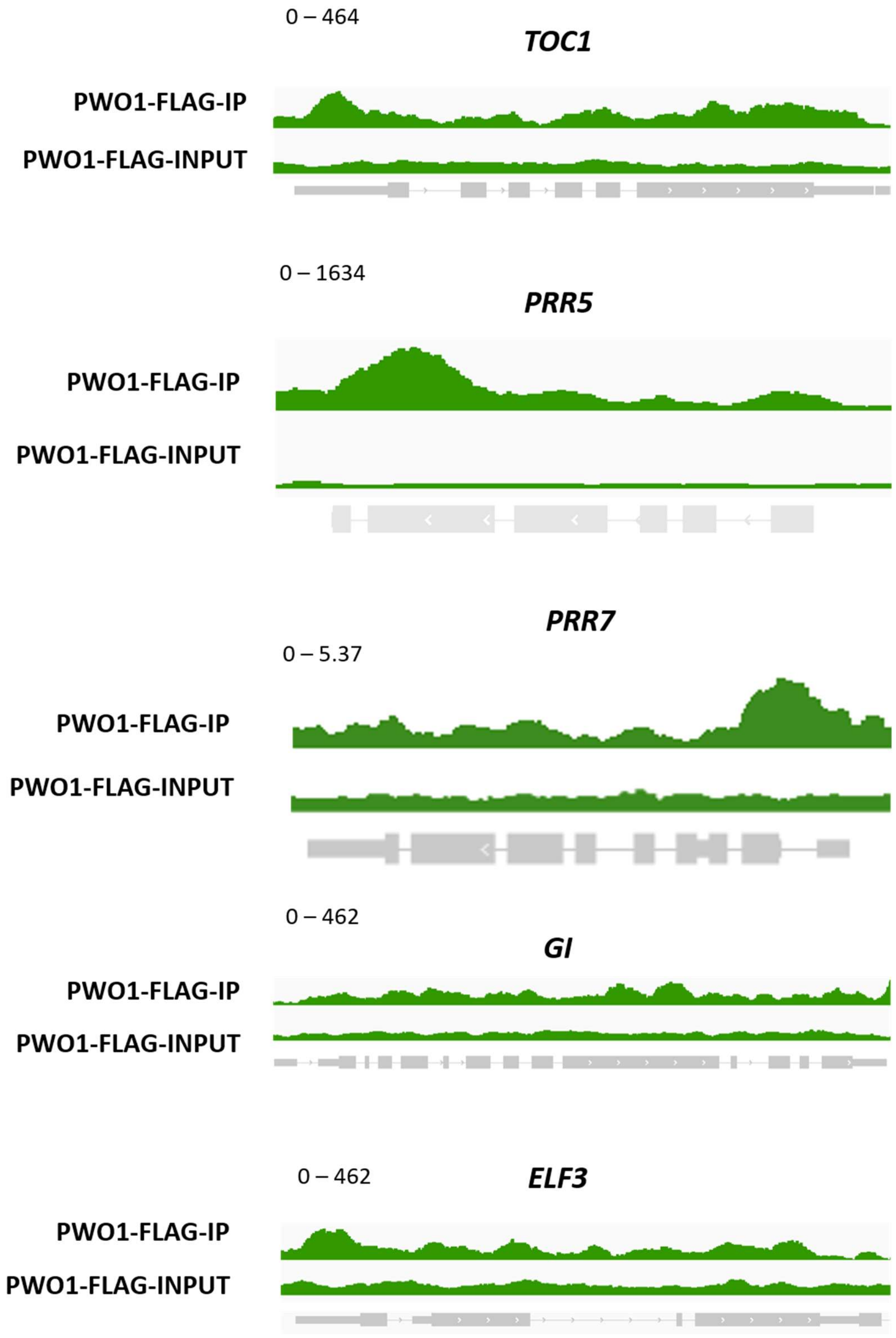


**Supplementary figure 3. PWO1-FLAG binding to circadian clock genes.** Integrative Genomic Viewer (IGV) snapshots of PWO1-FLAG binding at the *TOC1*, *PRR5*, *PRR7*, *GI*, and *ELF3* loci based on ChIP-seq results from Zheng *et al.* (2023).

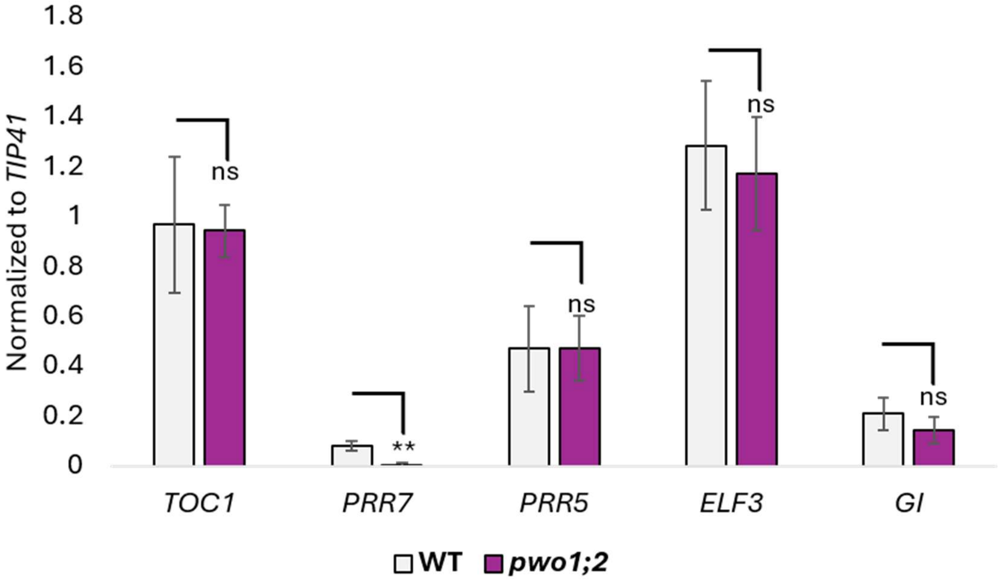


**Supplementary figure 4. Expression of selected PWO1 target genes in Col-0 (WT) and *pwo1;2* plants.** Expression levels of *TOC1, PRR7, PRR5, ELF3,* and *GI* analysed in 10-day-old seedlings grown under LD conditions. Samples were harvested at the end of LD (ZT16). *TIP41* was used as an internal control. At least 3 biological replicates were used. Student's t-test was used to calculate the statistical significance. Asterisks indicate significant differences between genotypes; *=P < 0.05, **=P < 0.01.

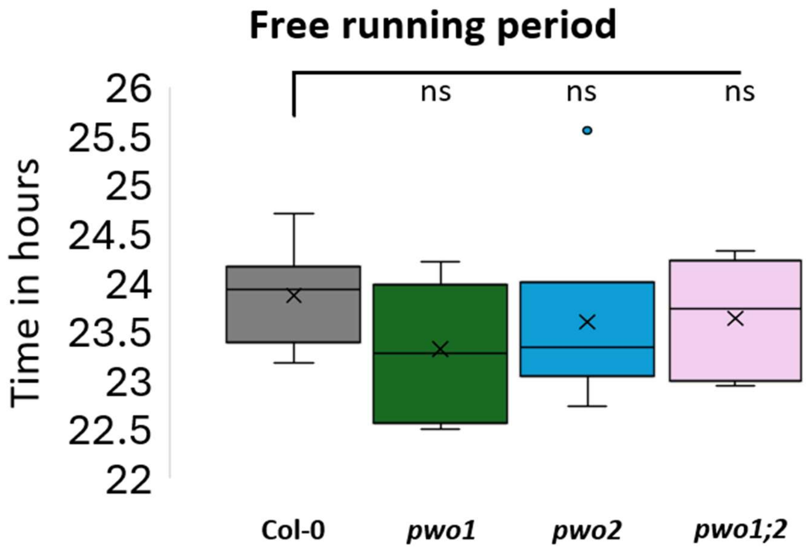


**Supplementary figure 5. Circadian period length based on rhythmic leaf movement in *pwo1, pwo2, pwo1;2* mutants compared to Col-0 (WT).** Box plot for showing the results for WT (n = 10), *pwo1* (n = 7), *pwo2* (n = 6), and *pwo1;2* (n = 5). Mean circadian period (hours) was similar across genotypes, ranging from 23.3 ± 0.68 h (*pwo1*) to 23.9 ± 0.45 h (WT). Across all parameters, no significant differences in central tendency were observed between *pwo1, pwo2, pwo1;2* mutants and WT following quality filtering via BioDare2, indicating broadly similar circadian timing under the conditions tested.

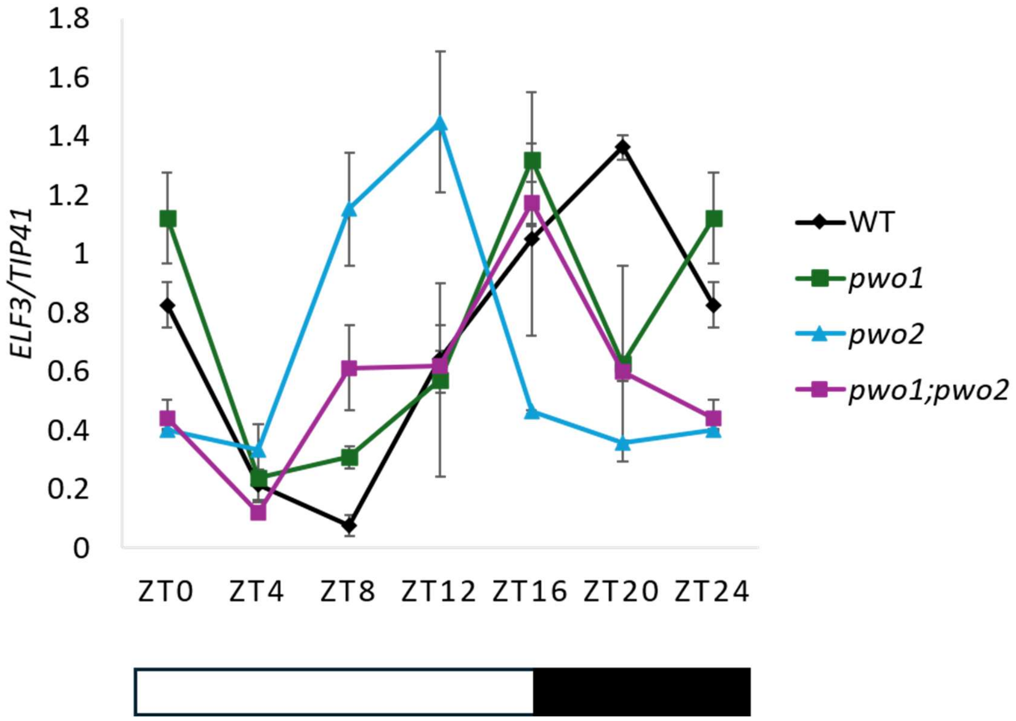


**Supplementary figure 6. Diel expression profiles of *ELF3* in *pwo1*, *pwo2*, *pwo1;2* compared to Col-0 (WT).** Expression levels of *ELF3* analysed in 10-day-old seedlings grown under LD conditions harvested every 4-hours. White and black bars represent day and night times respectively. *TIP41* was used as an internal control. At least 3 biological replicates were used.

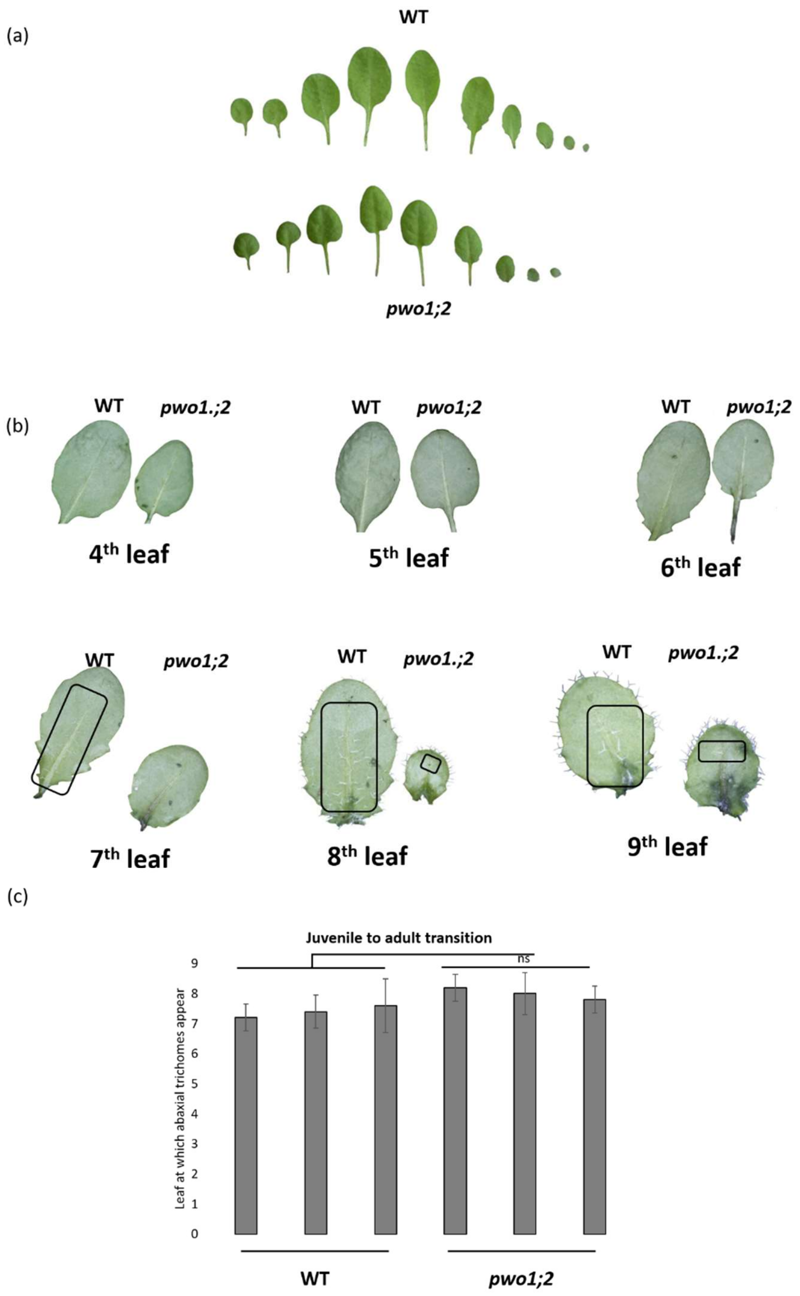


**Supplementary figure 7. Abaxial trichome analysis of Col-0 (WT) and *pwo1;2* under LD conditions.** (a-b) Representative leaf images and abaxial leaf surfaces of WT and *pwo1;2* plants grown under long-day (LD) conditions. Black square highlight regions containing abaxial trichomes. (c) The graph shows the leaf number at which abaxial trichomes first appeared, used as a marker of the juvenile-to-adult vegetative phase transition. 21-day-old plants were used for this analysis. Each bar represents data from 5 plants. Three biological replicates were used (n=15 in total).

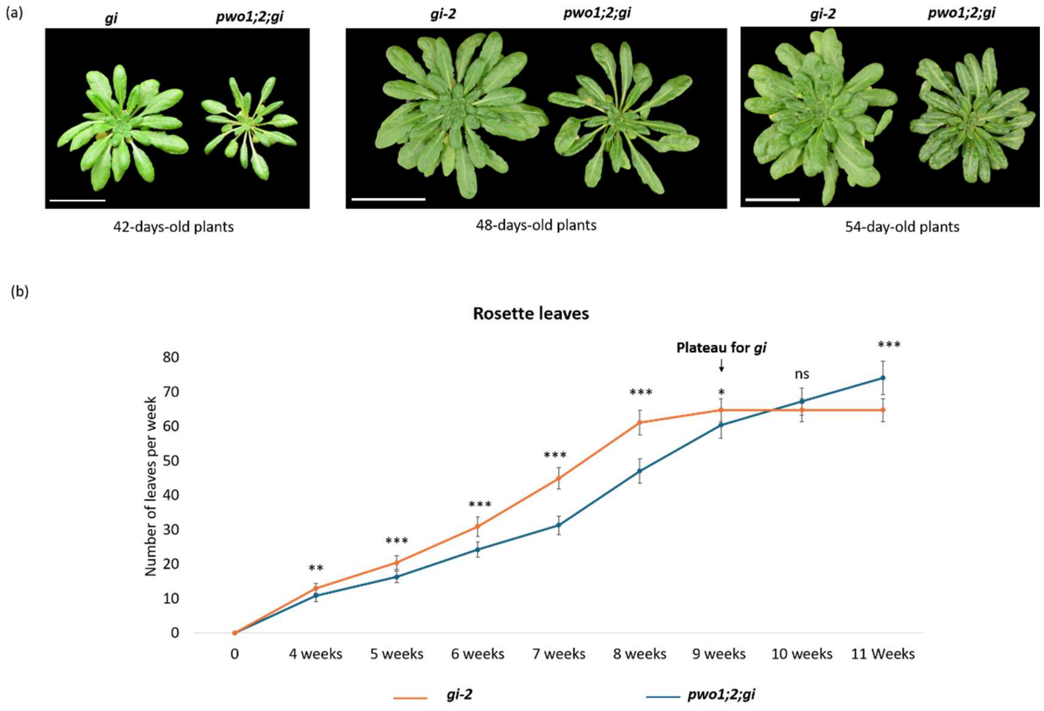


**Supplementary figure 8. Vegetative development and leaf production in *pwo;1;2;gi* compared to *gi* under LD conditions.** (a) Representative images of *gi* and *pwo1;2;gi* between 7-9 weeks. (b) Leaf number was recorded every week from 4 weeks onward to compare leaf production between *gi* and *pwo1;2;gi*. Error bars represent the SD. Student's t-test was used to calculate the statistical differences between the two genotypes (n=10). *=P < 0.05, **=P < 0.01, ***=P < 0.001. Scale bar = 5cm.

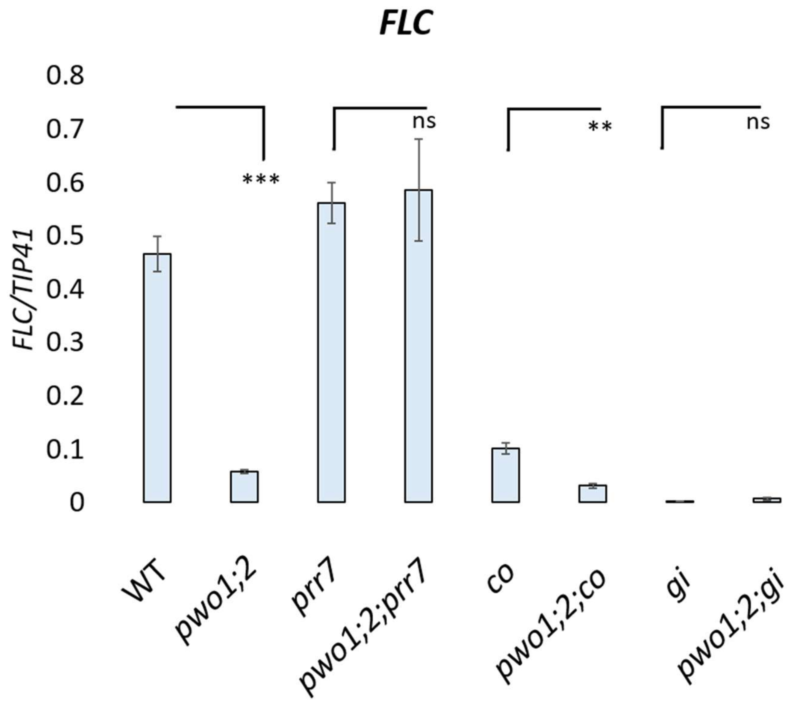


**Supplementary figure 9. *FLC* expression analysis 10-day-old seedlings in Col-0 and mutant genotypes.** Relative transcript levels of *FLC* in 10-day-old Col-0 *pwo1;2*, *prr7*, *pwo1;2;prr7*, *co*, *pwo1;2;co*, *gi*, and *pwo1;2;gi*. Samples were harvested at ZT16. Transcript levels were normalized to *TIP41*. Error bars represent the standard deviation from three biological replicates. Student`s t-test was used to calculate the significance level. Asterisks indicate significant differences between genotypes; *=P < 0.05, **=P < 0.01.
